# Force inference predicts local and tissue-scale stress patterns in epithelia

**DOI:** 10.1101/475012

**Authors:** W. Kong, O. Loison, P. Shivakumar, C. Collinet, P.F. Lenne, R. Clément

**Affiliations:** Aix Marseille University, CNRS, IBDM – Turing Center for Living Systems, Marseille, France

## Abstract

Morphogenesis relies on the active generation of forces, and the transmission of these forces to surrounding cells and tissues. Hence measuring forces directly in developing embryos is an essential task to study the mechanics of development. Among the experimental techniques that have emerged to measure forces in epithelial tissues, force inference is particularly appealing. Indeed it only requires a snapshot of the tissue, as it relies on the topology and geometry of cell contacts, assuming that forces are balanced at each vertex. However, establishing force inference as a reliable technique requires thorough validation in multiple conditions. Here we performed systematic comparisons of force inference with laser ablation experiments in three distinct *Drosophila* epithelia. We show that force inference accurately predicts single junction tensions, tension patterns in stereotyped groups of cells, and tissue-scale stress patterns, in wild type and mutant conditions. We emphasize its ability to capture the distribution of forces at different scales from a single image, which gives it a critical advantage over perturbative techniques such as laser ablation. Our results demonstrate that force inference is a reliable and efficient method to quantify the mechanics of epithelial tissues during morphogenesis.

## Introduction

During embryonic development, a small set of cell behaviors, including cell division, cell death and cell shape changes, lead to dramatic changes in tissue shapes. These events rely on forces generated at the cell scale, which build up and induce tissue scale movements, such as tissue elongation, tissue invagination or tissue closure (reviewed in (Heisenberg & Bellaïche 2013)). In epithelial morphogenesis of *Drosophila*, polarized contractile forces acting at cell junctions drive oriented cell intercalation and lead to convergent-extension (Bertet et al. 2004; Rauzi et al. 2008; Fernandez-Gonzalez et al. 2009). Stress generated locally can also propagate passively within surrounding cells and tissues as in the *Drosophila* posterior midgut, which largely contributes to elongating the adjacent germband upon invagination (Lye et al. 2015; Collinet et al. 2015). The tight genetic control of force generation leads to remarkably stereotyped shape changes, which is exemplified by the robustness of morphogenesis at the embryo scale. A consequence is that misregulation of force generation patterns leads to important morphogenetic defects. Interestingly, such robustness can hold at the scale of a few cells, as revealed by the strikingly regular cellular arrangements of the *Drosophila* retina (Hayashi & Carthew 2004).

A key step in understanding tissue morphogenesis is thus to establish reliable methods to assess the mechanical state of cells and tissues directly in the developing embryo. Evidently, measuring forces *in vivo* is not an easy task. A wide variety of techniques has recently been developed (for a review, see (Sugimura et al. 2016)), which include (but are not limited to) pipette aspiration (Guevorkian et al. 2010; Maitre et al. 2012), magnetic tweezers (Doubrovinski et al. 2017), laser cuts (Ma et al. 2009; Rauzi et al. 2008), or droplets injection (Serwane et al. 2017). All these techniques require to access the tissue of interest with a probe, and are therefore invasive and technically challenging. Optical tweezers have been used to perform non-invasive mechanical measurements at single junctions (Bambardekar et al. 2015; Clément et al. 2017), yet they only provide a small number of local measurements per embryo, and are thus difficult to implement to map the distribution of forces within a tissue. Force inference, which relies on the hypothesis that tensions equilibrate at each vertex, uses the geometry of cell contacts to infer a map of tensions and pressures from a tissue snapshot (Brodland et al. 2010; Chiou et al. 2012; Ishihara & Sugimura 2012; Brodland et al. 2014). Because it is fully non-invasive and does not require a specific experimental setup, it stands out as a simple and convenient method. As pointed out in a recent review (Sugimura et al. 2016), it is now crucial to cross-validate different measurement techniques in model systems in order to assess their robustness and reliability. Such cross-validation experiments require the combination of two or more techniques, and each of them being a technical challenge, cross-validation efforts remain rare in this rather new field of research.

Here, we investigate the accuracy of force inference using cross-validation with laser ablations experiments. Ishihara and co-workers combined force inference and annular laser cuts to show that force inference could predict coarse stress polarity averaged over the whole field of view in the *Drosophila* notum (Ishihara & Sugimura 2012; Ishihara et al. 2013). However a systematic, detailed cross-validation of force inference in different conditions and at different scales is chiefly missing, in particular for complex tension and stress patterns. To that end, we carried out our analysis in three different model tissues and at different spatial scales. We first validate our approach numerically on synthetic data. We then turn to the *Drosophila* notum, and study single junction tensions, showing that force inference correlates well with the recoil velocity of vertices following junctional laser cuts. We next turn to the *Drosophila* retinal ommatidia, and show that force inference adequately predicts tension patterns in these stereotyped groups of cells, in both wild type and mutant conditions. Finally, we show that force inference accurately predicts complex tissue-scale stress patterns revealed by line ablations in the wild type and mutant *Drosophila* germband, with unprecedented precision.

Altogether, our cross-validation study on different tissues demonstrates that force inference can be confidently used to assess the mechanical state of epithelial tissues. As accuracy increases with the level of coarse graining, we believe it is particularly well suited to determine complex stress patterns at the tissue scale during morphogenesis.

## Results

### 1 - Numerical validation of the inference methods

The inverse problem of force inference requires writing force balance equations at each vertex. To solve the set of equations, one thus needs to write and invert the corresponding matrix. A general difficulty is the indefiniteness caused by boundary conditions, where edges are connected to one vertex only. Because of Euler’s relation, the full inverse problem (one tension per edge and one pressure per cell) is indeed generally underdetermined (Ishihara et al. 2013). Different strategies can be adopted to handle the indefiniteness and yield a plausible set of tensions and pressures. First, one can assume that intracellular pressure is constant across the tissue. The problem then becomes overdetermined and can be solved by computing the pseudoinverse of the associated matrix (Chiou et al. 2012). As edges are rarely perfectly straight, suggesting pressure differences between cells, we chose to discard that assumption. Second, one can complement the contact angles measurements with the measurement of the radii of curvature between each pair of adjacent cells. Using Laplace’s law for each pair of adjacent cells, this provides another set of conditions that again leads to an overdetermined problem (Brodland et al. 2014). This is an ideal solution if edge curvatures can be accurately measured. Third, one can adopt a Bayesian approach, and incorporate statistical expectations for the system as a prior, for instance assuming a Gaussian distribution of tensions (Ishihara & Sugimura 2012). This is a good strategy when curvature measurements are difficult. In the Bayesian approach, pressure is written as a force acting on vertices (see supplements for details). In all cases, the inferred tensions are relative, as they are determined up to a multiplicative constant, while inferred pressures are determined up to an additive constant (hydrostatic pressure). A convenient convention is to scale tensions so that the average tension is 1, and to fix the reference average pressure to 0.

In this article, we used Bayesian inference (Ishihara & Sugimura 2012) for the notum and for the germband tissues (Figure 1A). The germband and the notum are rather close to hexagonal cell arrays, mostly composed of 6-sided cells. Proper curvature measurements are very challenging in these tissues, as 1/ curvatures are usually rather small and 2/ upon segmentation, cell edges often have the shape of a very open S, making it impossible to properly determine curvature (Figure S1). In contrast, we used Laplace inference (Brodland et al. 2014) for the ommatidia (Figure 1F). Indeed, ommatidia are stereotyped units composed of 6 cells with highly stereotyped shapes and high curvature, which allow averaging and therefore much easier and reliable measurements of curvature.

**Figure 1.**
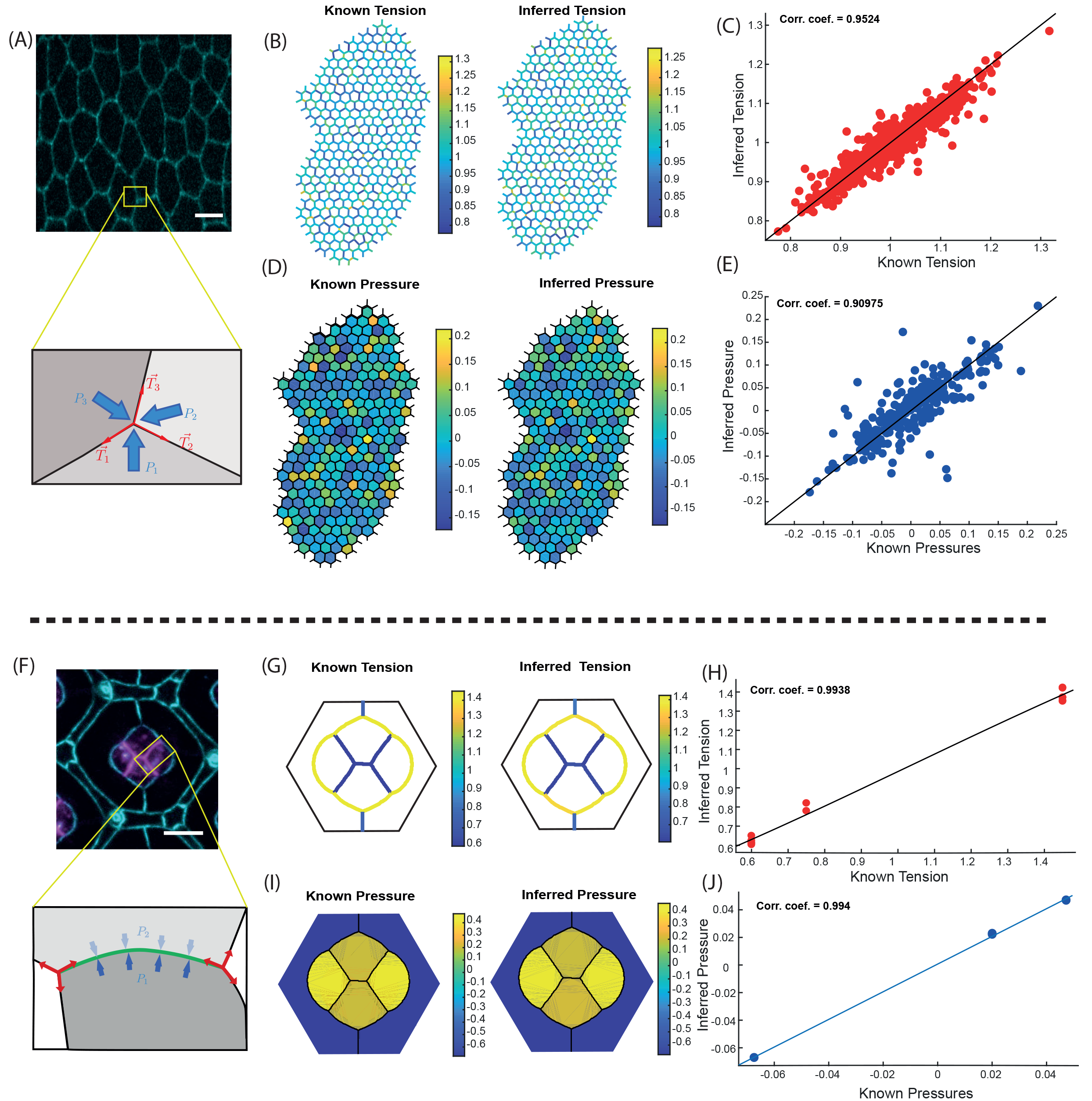
Validation of force inference on synthetic data. **(A)** Scheme of the force balance in the Bayesian force inference method. Tensions and pressures act as forces directly applied to vertices, and the force balance is written at each vertex. Sample image: Drosophila notum around 22h after pupa formation. Scale bar: 5μm. **(B)** Known (left) vs inferred (right) tension map in a tissue simulated with Surface Evolver (246 cells, 873 edges). **(C)** Inferred tensions plotted against their known value. **(D)** Known (left) vs inferred (right) pressure map. **(E)** Inferred pressures plotted against their known value. **(F)** Scheme of the force balance in the Laplace force inference method. Tensions act as forces applied to vertices tangent to the edges, and pressures are related to edge tension and curvature by Laplace’s law. Sample image: Drosophila ommatidium 41h after pupa formation. Scale bar: 5μm. **(G)** Known (left) vs inferred (right) tension map in an ommatidium simulated with Surface Evolver. **(H)** Inferred tensions plotted against their known value. **(I)** Known (left) vs inferred (right) pressure map. **(J)** Inferred pressures plotted against their known value.

Before applying our force inference custom codes to living tissues, we first checked that they were properly implemented using *in silico* tissues generated with the Surface Evolver software (Brakke 1992). Surface Evolver uses energy minimization to drive a system governed by custom line/surface energies to equilibrium.

To validate our Bayesian inference code, we first computed a hexagonal array of cells. We then assigned tensions and pressures to edges and cells, respectively. Tensions have a Gaussian distribution centered around 1. We let the system reach equilibrium by minimizing the energy of a classical vertex model (Farhadifar et al. 2007), and finally inferred tensions and pressures from the resulting geometry. Thus we could directly compare inferred tensions and pressures to known tensions and pressures. Bayesian inference performs very well, as shown by comparisons between the true versus inferred tension and pressure maps (Figures 1B, 1C). The correlation is excellent for both tensions and pressures, with a Pearson’s correlation coefficient above 0.9 (Figure 1D, 1E).

We used a similar validation approach for our Laplace inference code, this time using simulations of groups of cells mimicking *Drosophila* ommatidia, which we use as a model system later in this study. In silico ommatidia were also generated using a classic vertex model in Surface Evolver. Again, we find an excellent agreement between simulations and force inference, as shown by the tension and pressure maps (Figures 1G, 1H). Despite the limited size of the system, which only has 6 cells and 13 edges, the correlation remains excellent for both tensions and pressures (Figures 1I, 1J).

### 2 Single junction tensions in the pupal notum

The most straightforward experimental verification of force inference accuracy is to directly compare tensions inferred in single junctions to measurements obtained from single junction laser ablation, which is the most used experimental technique to evaluate tensions. In laser ablation experiments, a tightly focused laser disrupts the molecular structures that support tension in a targeted junction. The recoil velocity following a single junction laser cut is proportional to the tension of the targeted junction just prior to the cut (Rauzi & Lenne 2015). Indeed, upon release, tension is only balanced by fluid friction. To compare force inference to laser cuts, we turned to a rather regularly organized epithelium, the pupal notum of *Drosophila* around 21h after pupa formation (Figure 2A). Tension variations at this stage are not expected to be particularly oriented, as revealed by annular laser cuts (Bonnet et al. 2012). Hence they are essentially random fluctuations that cause the system to slightly deviate from a regular hexagonal array. Because force inference provides relative estimates, it is always delicate to compare tensions estimated from separate images. We thus hypothesized that the average tension was always the same in all of our images (normalized to 1). To moderate the influence of this assumption, for each field of view where force inference was performed, we did several (3 to 5) laser cuts, sufficiently spaced so as not to influence each other (Figure 2A, 2B, S2). For direct comparison, force inference was always computed in images taken just prior to the laser cuts. We then compared the inferred tensions to the initial recoil velocities of the considered junctions, measured by fitting the onset of the opening (Figure 2C). We found a fairly good correlation coefficient of about 0.6 between opening velocities and inferred tensions (Figure 2D). The discrepancy can arise from numerous sources: the intrinsic hypotheses of force inference, but also the errors made on velocity measurements, mainly the assumptions that tension is solely balanced by pure fluid friction and that fluid friction is homogeneous in the tissue. The correlation found despite these limiting factors suggests that both methods can provide reliable results. Clearly, single junction tension measurements are also prone to more errors than tensions averaged over groups of junctions. Such groups can be based on position (coarse graining), orientation (to detect polarity), or biological identity. We therefore questioned next whether force inference could detect tension gradations between different, stereotyped groups of junctions.

**Figure 2.**
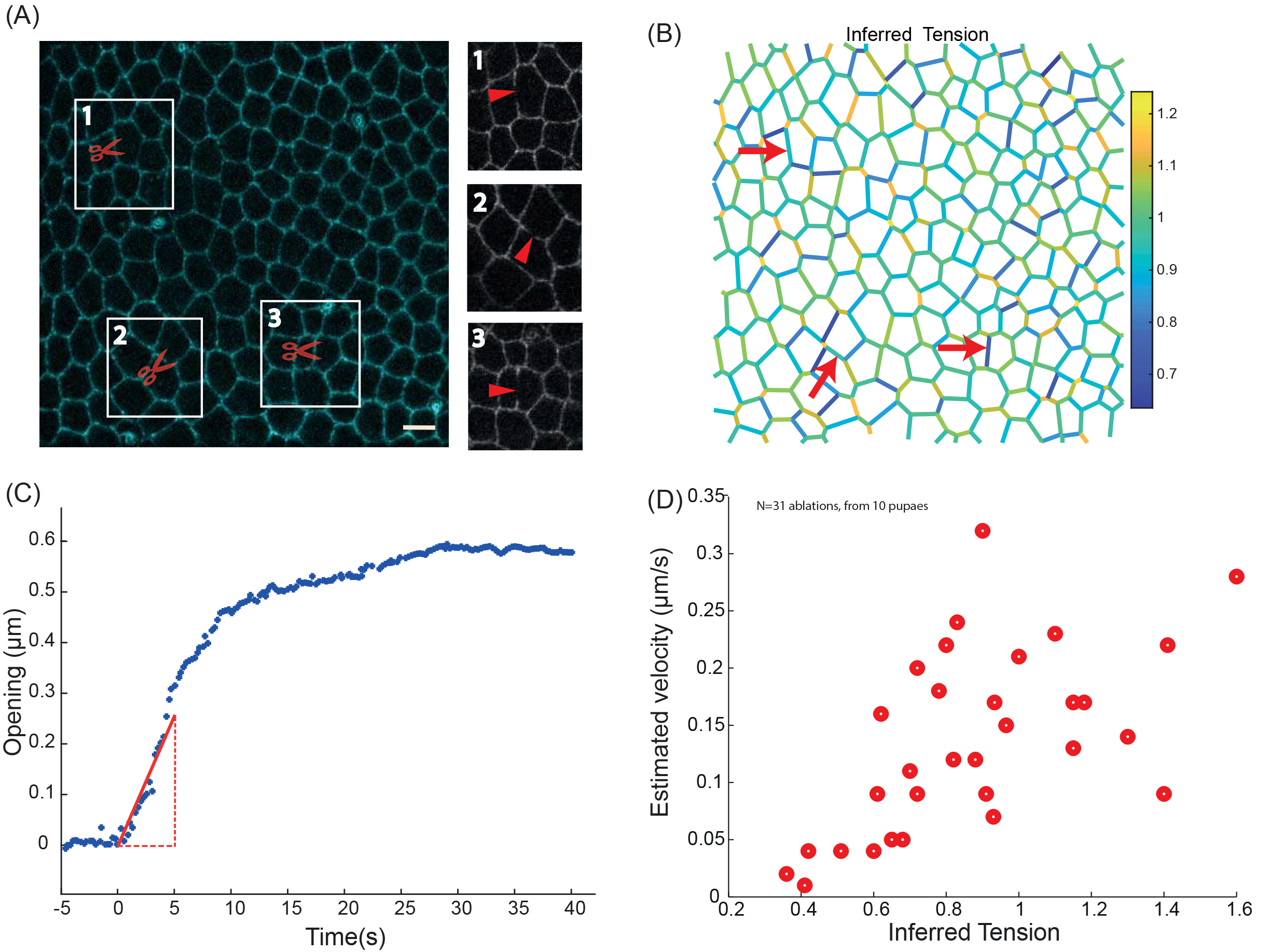
Force inference at the single junction scale in the Drosophila notum. **(A)** Subregion of the Drosophila notum 21h after pupa formation. Scissors show ablation spots where recoil velocities will be measured. Insets show post-ablation snapshots of the considered junctions. Scale bar: 5μm. **(B)** Inferred tension map of the tissue region in (A) before the ablations. Red arrows indicate the location of ablations, where inferred tensions are extracted and compared to experimental recoil velocities. **(C)** Opening dynamics and initial recoil velocity. The red line show a linear fit of the first 5 seconds, which is used to determine the initial recoil velocity. **(D)** Opening velocity plotted against inferred tension for N=31 laser cuts performed in 10 pupae. Pearson’s correlation coefficient is 0.59. Spearman’s correlation coefficient is 0.63.

### 3 Stereotyped tension patterns in wild type and mutant ommatidia

Stereotyped patterns of differential tension between subgroups of cells can drive robust geometric organization of multicellular structures. We wanted to assess whether force inference could detect such patterns of tensions. To achieve this, we turned to the retina of *Drosophila*, composed of highly stereotyped groups of cells called ommatidia (Figure 3A). Previous studies showed that cone cell shapes and arrangements in ommatidia are determined by differential tensions (Hayashi & Carthew 2004; Käfer et al. 2007; Hilgenfeldt et al. 2008; Chan et al. 2017). These tensions were shown to be determined by the amounts of Myo-II and E- and N- cadherins recruited at the considered junctions (Chan et al. 2017). These amounts were in turn shown to be determined by the “identity” of junctions, that is, by the types of cadherins expressed in the two contacting cells (Chan et al. 2017). Based on these previous results, we categorized junctions according to the cadherins expressed in the contacting cells. EN|EN junctions correspond to homotypic junctions separating two cells that both express E- and N-Cadherin. E|E junctions correspond to homotypic junctions separating two cells that both express E-Cadherin only. EN|E junctions correspond to heterotypic junctions separating a cell expressing E-Cadherin only from a cell expressing both E and N-Cadherin. These three types of junctions coexist in a wild type ommatidium (Figure 3B).

**Figure 3.**
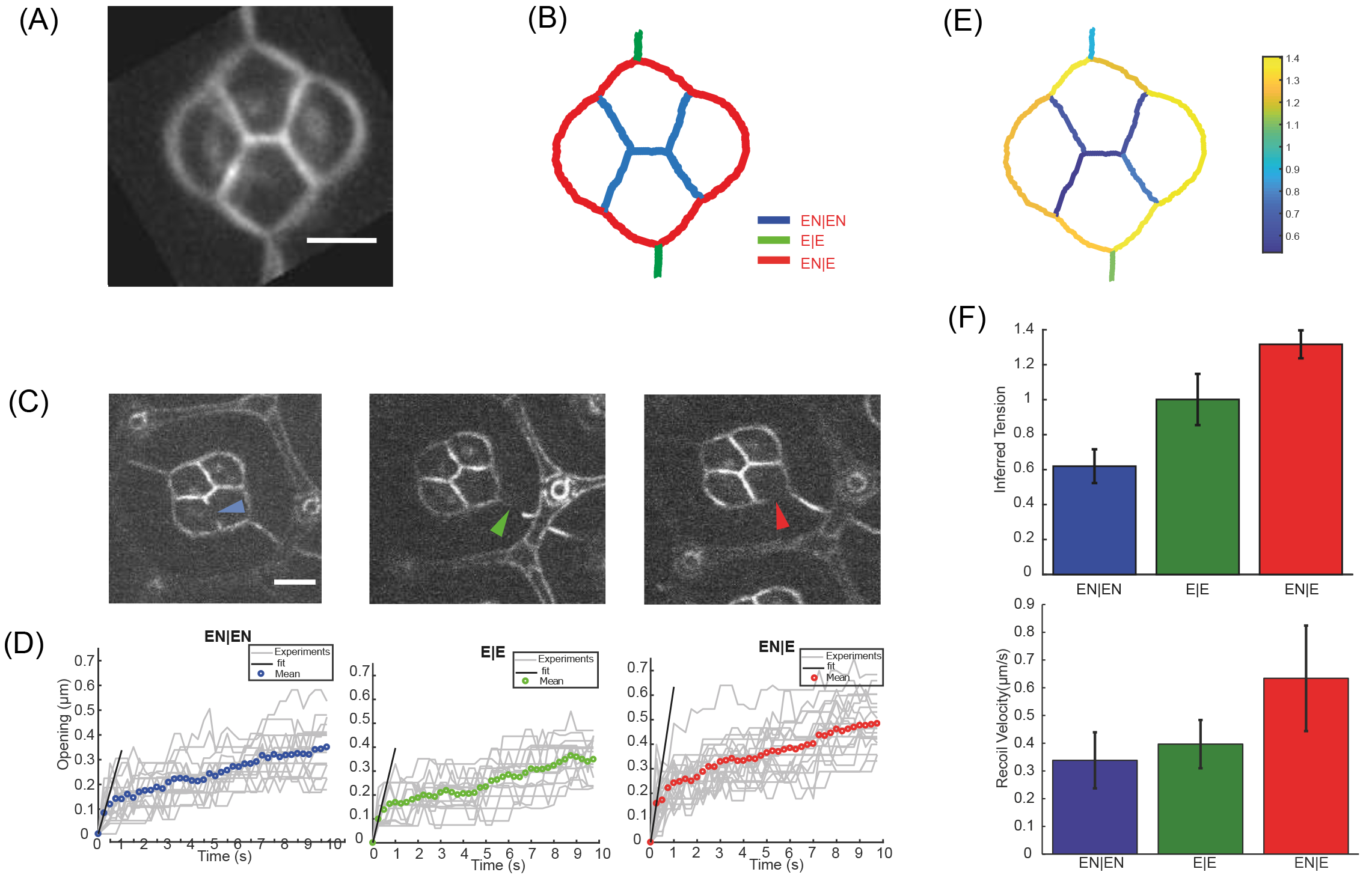
Force inference in the WT Drosophila retina. **(A)** The four cone cells of a Drosophila ommatidium. The image results from an average over N=51 ommatidia, which we use for force inference. Scale bar: 5μm. **(B)** Segmented version of (A), and nomenclature of the junction types: EN|EN junctions are depicted in blue, E|E in green, and and EN|E in red. **(C)** Snapshots of ablation experiments performed for each type of junction. **(D)** Averaged opening dynamics for each type of junction (EN|EN: N=19, E|E: N=16, EN|E: N=22). Individual opening curves are shown in light gray. **(E)** Tension map inferred from the segmented average ommatidium. **(F)** Top panel: average tension inferred for each junction type. Error bars: Standard deviation. Bottom panel: Average opening velocity measured for each junction. Error bars: standard error on velocity (computed from the standard error on position).

We computed the averaged opening dynamics following laser cuts for each type of junctions, and extracted the corresponding initial recoil velocity (Figure 3C, 3D). As previously demonstrated (Chan et al. 2017), this revealed a gradation of tensions according to junction type. Homotypic EN|EN junctions have the lowest tensions, E|E junctions have intermediate tensions, and heterotypic EN|E junctions have the highest tensions (Figure 3F). Note that tensions are directly related to the amounts of Myo-II present at these junctions (Chan et al. 2017). To perform force inference in this system, we averaged ommatidia geometry over N=51 ommatidia, and segmented the resulting image (Figure 3A, 3B). As stated earlier, high and stereotyped curvatures in this system make it possible to properly measure the tangents and radii of curvature required for Laplace inference. We found that inference accurately predicts the pattern of tensions and its gradation among the three types of junctions (Figure 3E, 3F). Note that cell pressures can also be computed (Figure S3).

We then turned to the analysis of mosaic experiments in which a fraction of cells do not express N-Cadherin (Chan et al. 2017). Since the mutation affects random cells in the tissue, such experiments generate a variety of configurations, in which a subgroup of one or more cone cells are affected by the mutation (Figure 4A). Interestingly, this effectively modifies the spatial distribution of the junction types in the ommatidia, since the junction type is determined by which cadherins are expressed by the contacting cells (Figure 4B). To test whether force inference could still detect tension gradation in these modified conditions, we applied force inference to 5 different configurations of ommatidia (Figure 4C). Note that, due to the stochastic generation of these configurations, inference is performed on a single ommatidium for each configuration, whereas an average over many ommatidia was used for the wild type condition. Strikingly, the gradation of tensions identified in the wild type condition is systematically detected by force inference in the various mutant configurations (Figure 4D). This suggests that tensions are indeed determined by the combination of cadherins expressed by adjacent cells, through adhesion strength but also Myo-II level (Chan et al. 2017). Inference results are also consistent with laser cuts averaged over all mutant configurations for each junction type (Figure S4).

**Figure 4.**
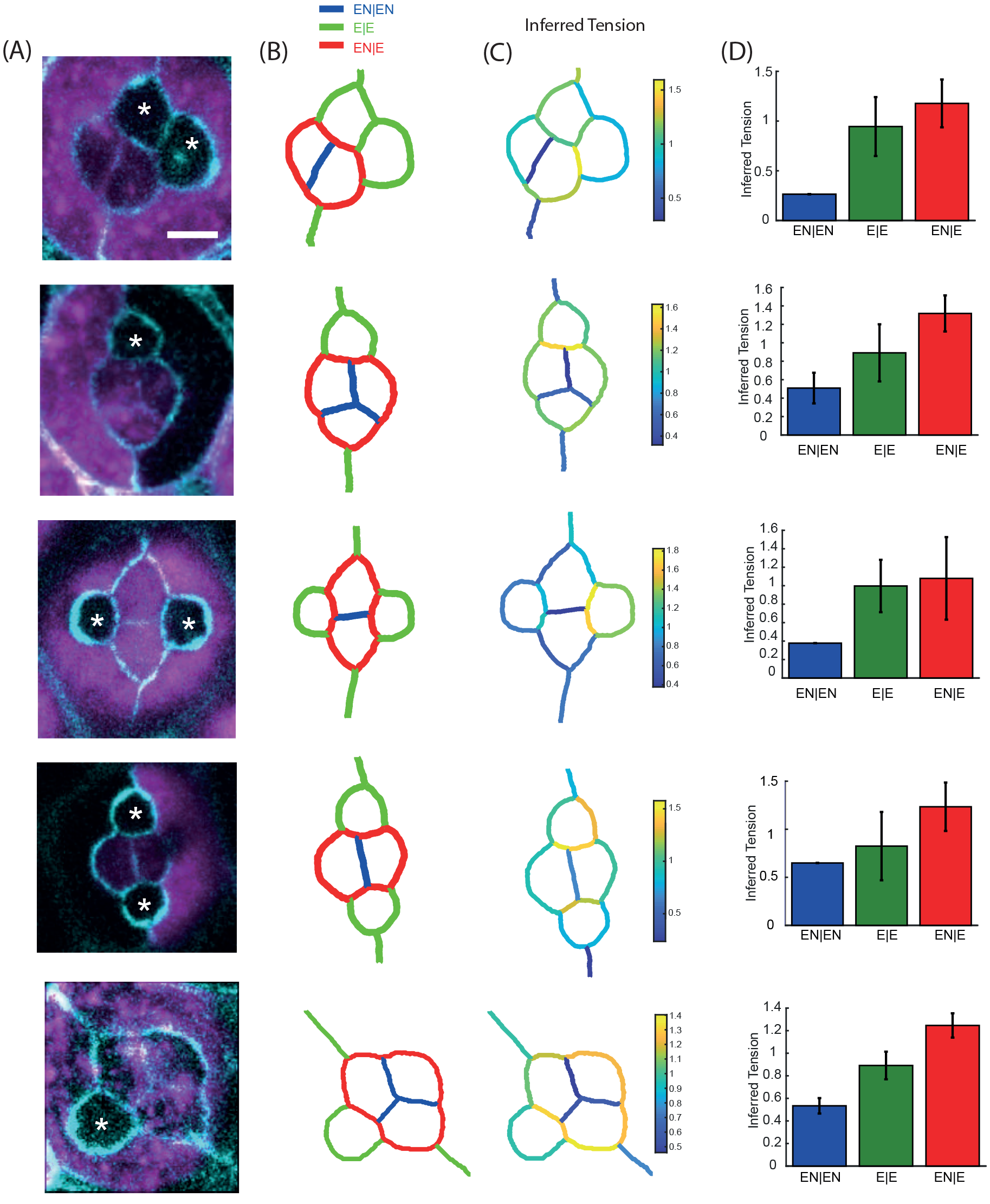
Force inference in mutant Drosophila retina. **(A)** Five different mutant configurations generated from the mosaic experiments. WT cells are in purple. Untagged cells do not express N-Cad. This only affects cone cells, as surrounding cells do not express N-Cad. Starred cells are thus N-Cad negative cells. Scale bar: 5μm. **(B)** Pattern of junction types for each configuration. **(C)** Map of inferred tension for each configuration. **(D)** Average tension for each junction type in each configuration. Error bars: standard deviation.

Overall, the results obtained in the retina suggest that force inference can robustly detect tension patterns in stereotyped units of a few cells. This led us to investigate the ability of this technique to detect stress patterns at the scale of hundreds of cells, relevant to many morphogenetic processes.

### 4 Stress patterns in the wild type and mutant germband

Force inference, by coarse graining tensions and pressures at the appropriate scale, can be used to build a map of the stress tensor (Batchelor 1970; Ishihara & Sugimura 2012). Very promising results were obtained using this approach to determine the complex stress pattern of the entire notum (Guirao et al. 2015). However comparison to experimental stress measurements was only performed at a rather coarse level, looking at the overall anisotropy of the whole field of view, by binning tensions and pressures on the whole sample (Ishihara et al. 2013). To determine whether this approach could predict complex stress patterns in more details, we turned to a mechanically well-characterized tissue, the embryonic germband of *Drosophila*, that is known to display stress polarity induced by Myo-II polarity, but also a stress gradient along the antero-posterior (AP) axis, due to the movement of the posterior midgut pulling on the tissue (Collinet et al. 2015).

In the wild type germband, the polarized recruitment of MyoII at dorso-ventral (DV) junctions is known to polarize stress and induce polarized intercalation of cells. In addition, the posterior midgut, which undergoes rotation and invagination, has been shown to pull on the germband along the AP axis, inducing an additional gradient of stress along this axis (Figure 5A, left panel). This is illustrated by the opening velocities measured following large AP- or DV-oriented line cuts performed in the anterior, middle and posterior regions of the germband (Figures 5B, left panel). In the anterior region, away from the posterior midgut, stress is dominated by Myo-II polarity and is strongly polarized along the DV axis. In the middle region, as we get closer to the posterior, stress along the AP axis increases, but remains smaller than stress along the DV axis. In the posterior region, stress along the AP axis becomes even larger due to the proximity to the posterior midgut, and stress along the AP and DV axes become comparable, so that stress polarity is lost. We performed force inference on the germband during this process. The tension map suggests that tensions are indeed higher along the DV axis (Figure 5C, left panel). To obtain a better representation of polarity, we computed the stress tensor, binning over square subregions of typically 8-10 cells. We then plotted its principal directions in each subregion, using the corresponding eigenvalues as amplitudes (Figure 5D, left panel). The results are fully consistent with the laser cut experiments. In the anterior region, stress is largely polarized along the DV axis. As we get closer to the posterior, stress along the AP axis gradually increases, so that in the posterior region, DV polarity is gradually lost. Interestingly, the gradual decrease of stress along the AP axis suggests that dissipation occurs across the tissue, over a length scale of typically 100 μm (the hydrodynamic length). To further test force inference ability to detect stress patterns, we used mutant conditions in which posterior midgut invagination (Torso −/−) or Myo-II polarity (Eve RNAi) are selectively impaired. In the absence of posterior midgut invagination, posterior pulling forces are abolished (Figure 5A, middle panel), and the stress pattern is mostly determined by Myo-II polarity, with an important DV stress polarity from anterior to posterior (Figure 5B, middle panel). This is fully recapitulated by the force inference approach (Figures 5C, 5D, middle panels). In contrast, in the absence of Myo-II polarization, the stress pattern is mostly determined by the posterior forces (Figure 5A, right panel). Laser cuts show that stress along the DV axis is reduced across all the tissue, while the gradual increase of stress along the AP axis from anterior to posterior is maintained (Figure 5B, right panel). The stress pattern is again fully recapitulated by the force inference approach (Figures 5C, 5D, right panel).

**Figure 5.**
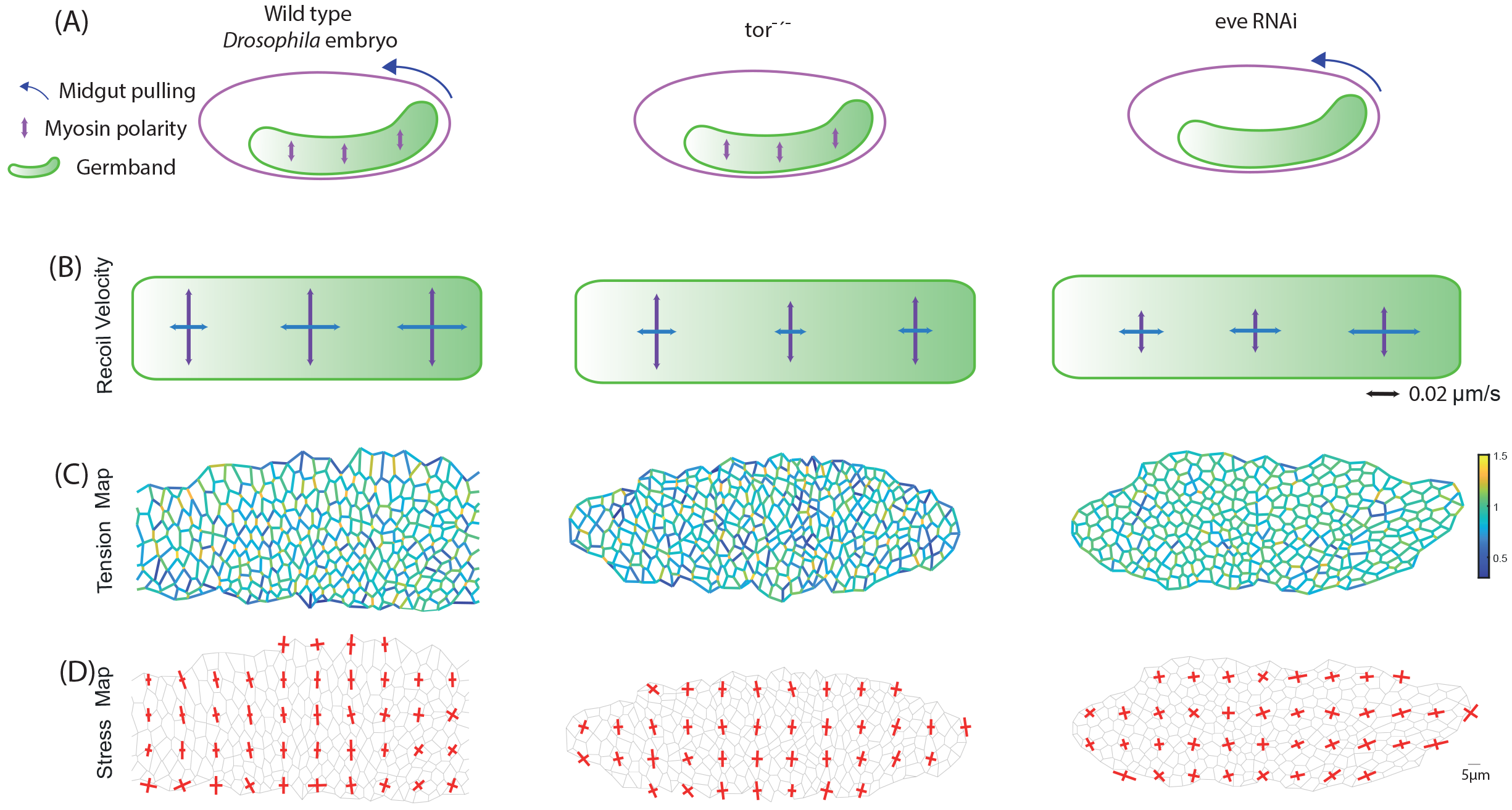
Tissue scale force inference in the Drosophila germband. **(A)** Scheme of stress sources in the germband. In the WT condition (left), Myo-II polarity generates stress along the DV axis, and posterior midgut invagination pulls on the germband from on its posterior side. In the Tor−/− condition (middle), posterior midgut invagination is abolished, and Myo-II polarity is preserved. In the Eve RNAi condition (right), posterior midgut invagination is preserved, and Myo-II polarity is lost. **(B)** Recoil velocities measured with PIV for each condition in the anterior, middle and posterior regions of the germband. Vertical arrows correspond to opening velocities along the DV axis (cuts along the AP axis), and horizontal arrows correspond to opening velocities along the AP axis (cuts along the DV axis). **(C)** Inferred tension map in each condition. **(D)** Inferred stress map computed with Batchelor’s formula from the inferred tensions and pressures.

Interestingly, derivation of an approximate stress tensor can be done without solving the inverse problem, if one assumes that all tensions and pressures are homogeneous. This actually yields a surprisingly good approximation (Figure S5), which is solely based on the contribution of cell shape anisotropies detected by segmentation. An advantage is that it is much easier to implement and faster to compute, since solving the inverse problem can be computationally demanding in very large systems. In that context, one could imagine that even segmentation might not be required, if cell shape anisotropies can be detected from proper spectral analysis of the tissue image. This could certainly be an interesting option for tissues in which segmentation is challenging due to imaging difficulties.

Altogether, the analyses of normal and impaired germband extension show that force inference can precisely recapitulate stress patterns across a dynamic epithelium undergoing morphogenetic movements. Force inference is also much faster than laser ablations, and the spatial resolution of the estimated stress is much higher.

## Discussion

Advantages of force inference include that it is fully non-invasive, much easier to perform than perturbative experimental measurements, and does not require assumptions on the origin of forces involved or on the tissue rheology. Whether the criteria required to confidently use force inference are generally met in classic epithelial model systems was still unclear. Here, we conducted a thorough cross-validation in epithelia at different developmental stages and with very different geometries and dynamics. Our results demonstrate that force inference can be reliably used to analyze the mechanical state of epithelia, from the cell scale up to the tissue scale.

Force inference allows fairly good estimates of tension, even at the single junction scale. By providing a large set of measurements from single images, force inference can be an asset to search for correlations between tensions and protein distributions with good statistics (Kale et al. 2018). Clearly, averaging over groups of junctions of interest, or coarse-graining tensions and pressures into a binned stress tensor, significantly improves the reliability of the pattern detected. For the germband, a highly dynamic morphogenetic tissue with a complex stress pattern, the advantages of using inference were obvious compared to laser cuts, which can be painfully long to perform as they require averaging over many animals. Inference over a single, well segmented germband not only recapitulated the ablation findings but also allowed a more precise characterization of the stress pattern. This shows that force inference is particularly well-suited to determine stress patterns at the tissue scale during morphogenetic events, as previously done by Guirao and coworkers (Guirao et al. 2015). Hence it can also be an asset to study stress propagation and tissue rheology during morphogenesis, as tissue- or even animal-scale stress patterns and tissue flows can be established by active forces generated locally (Dicko et al. 2017). A drawback of force inference is that it only provides relative measurements, which can make it difficult to compare different animals or conditions in the absence of a reference. When required, this difficulty can be overcome either by calibration of a reference tension with another technique, or more simply by an assumption of equal mean tension when reasonable (same conditions in different animals, for instance).

## Author Contributions

WK conducted the laser cut experiments in the notum and analyzed the data. PS conducted the laser cut experiments in the ommatidia, WK and PS analyzed the data. CC conducted the laser cut experiments in the germband and analyzed the data. RC, WK, and OL implemented the force inference codes and their verification with Surface Evolver. WK conducted the force inference analysis in the notum and in the retina. RC conducted the force inference analysis in the germband. RC and PFL designed the project. RC, WK and PFL discussed the data. RC and WK wrote the paper. All authors commented on it.

## Acknowledgements

We thank Eunice Chan for the images of mutant ommatidia, Claire Chardès for her assistance with the laser ablation setup, and S. Ishihara and K. Sugimura for their helpful comments about the Matlab implementation of Bayesian force inference and QR decomposition using SuiteSparse. We thank members of Lenne and Lecuit groups for discussions throughout the course of this project and for providing a stimulating environment. WK is supported by Ph.D. fellowship from the LabEx INFORM (ANR-11-LABX-0054) and of the A*MIDEX project (ANR-11-IDEX-0001-02), funded by the “Investissements d’Avenir” French Government program. The project was in part funded by the ANR grant ANR-17-CE13-0032. We acknowledge France-BioImaging infrastructure supported by the French National Research Agency (ANR-10-INBS-04-01, «Investments for the future»).

## Methods

### 1 Segmentation

We used the Tissue Analyzer plugin for FIJI to segment our images (Aigouy et al. 2010). The segmented data (vertices, edges, connectivity) was then passed to Matlab and used for force inference.

### 2 Bayesian force inference

Bayesian force inference was implemented in a custom Matlab script. The mathematical formulation of the method was first introduced by Ishihara and coworkers (Ishihara & Sugimura 2012). The curvature of edges is not taken into account to solve the problem, so that tensions are assumed to be directed along cords joining vertices. Tensions and pressures are thus determined simultaneously, as both tensions and pressures are written as forces acting directly on vertices. The problem is therefore underdetermined. A Gaussian prior on tension distribution is used to overcome the underdetermination. We used SuiteSparse to perform QR decompositions in Matlab (Davis 2011).

The stress tensor (Figures 5D) is determined using Batchelor’s formula (Batchelor 1970; Ishihara & Sugimura 2012):

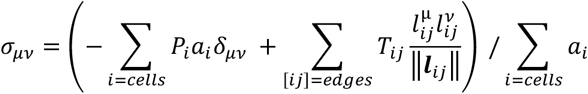

where *a*_*i*_ is the area of cell *i*, *P*_*i*_ its pressure, *δ* is Kronecker’s symbol, *T*_*ij*_ is the tension of the edge [*ij*] separating cells *i* and *j*, and ***l***_***ij***_ the vector connecting the two vertices of edge [*ij*]. Red bars show principal directions of *σ*, and their length is proportional to the corresponding eigenvalues. Note that it is up to the user to decide the appropriate level of coarse-graining. Stress can be computed separately for each cell, or averaged over subregions of any desired size (here, 8-10 cells).

### 3 Laplace force inference

Laplace force inference was implemented in a custom Matlab script. The mathematical formulation of the method used was introduced by Broadland and coworkers (Brodland et al. 2014). First, tensions are determined separately by measuring the tangents at each vertex and solving the force balance system (which is then independent of pressures and thus overdetermined). To determine the tangents we performed linear fits of the first 7 pixels of each edge (for each vertex). Next, the curvature of each junction is determined using Taubin circle fitting method (Taubin 1991). Pressures can then be determined using Laplace’s law for each pair of adjacent cells, which again yields an overdetermined system. Note that pressure determination is not crucial to our analysis of ommatidia, since we have no experimental data concerning cell pressures.

### 4 Tissue simulations (Surface Evolver)

The synthetic tissue data was generated using Surface Evolver v2.7 (Brakke 1992). Surface Evolver evolves the given surface towards its minimal energy configuration by a gradient descent method. In our case we use a classical vertex model energy of the form (Farhadifar et al. 2007; Chan et al. 2017):

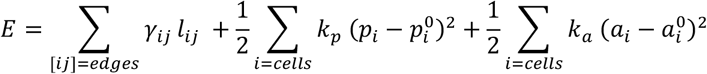

where *p*_*i*_ and *a*_*i*_ are the perimeter and area of cell *i*, and 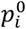 and 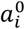 its target perimeter and target area. *k*_*p*_ and *k*_*a*_ are the strengths associated to the perimeter and area constraints, respectively. *γ*_*ij*_ is the line tension in edge [*ij*], and *l*_*ij*_ its length.

The pressure in cell *i* (“known” pressure) is then given by 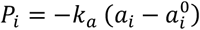, and the total tension of edge [*ij*] (“known” tension) is given by 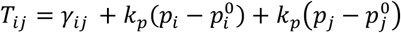

In the tissue simulation (Figure 1B), the target area is set to 0.86603 and *k*_*a*_ is set to 2. For the sake of simplicity the target perimeters are all set to 0. *k*_*p*_ is set to 0.15. Line tensions *γ*_*ij*_ are randomly assigned from a Gaussian distribution (mean=1, std=1/6) prior to equilibration.

In the ommatidium simulation (Figure 1G), target area are set to 5.5 for top-bottom cone cells, 5 for right-left cone cells and 40.49 for surrounding cells. *k*_*a*_ is set to 38.2 and *k*_*p*_ is set to 0. Tensions *T*_*ij*_ = *γ*_*ij*_ are directly set to 0.75 for E|E junctions, 0.6 for EN|EN junctions, and 1.45 for EN|E junctions.

Note that whether these values are realistic or not is of little concern, since their only purpose is to demonstrate the accuracy of the force inference codes.

### 5 Flies

In the notum, Ecad:GFP/Sqh:MCherry flies were used.

In *Drosophila* ommatidia, E-CAD:GFP; N-CAD:mkate2 flies were used (Chan et al. 2017). Mosaic experiments were also described in a previous paper (Chan et al. 2017).

In the germband, a; E-cad::GFP^KI^ fly line was used as wild type, embryos from a; tor^4^, E-cad::GFP^KIn^ were used as Torso −/− and dsRNAs against even-skipped injected in embryos form; E-cad::GFP^KI^ flies as previously described (Bertet et al. 2004; Collinet et al. 2015) to obtain eve loss-of-function embryos.

### 6 Laser ablations

All the laser ablation experiments were performed on a previously described setup (Rauzi et al. 2008). Junction cuts in the retina were described in a previous article (Chan et al. 2017). Line cuts in the germband were also described in a previous article (Collinet et al. 2015).

### 7 Measurement of recoil velocities

In the notum, we used kymographs along lines parallel to the severed junctions to automatically track the movement of vertices. Kymographs were then oversampled and treated with a Gaussian filter to avoid pixelation effects in vertex detection. Vertices positions were determined at each time point with a Gaussian fit of the intensity. Despite these efforts, the data can still be quite noisy. In addition, we needed to fit separately each single opening curve, without the possibility to average over several junctions as we wanted to do single junction comparisons with force inference. Thus to determine the initial recoil velocity we had to perform a linear fit on the first 5 seconds. This was determined empirically, as too short fitting times are very much affected by noise, and too long fitting times yield poor estimates as opening is rather exponential or biexponential than linear.

In the retina, automated detection with kymographs could not be used, due to smaller cells, edge curvatures, and higher signal loss following ablations. Hence, we used a manual tracking approach of the vertices, using FIJI. In this case, opening curves could be averaged over several junctions, which yielded much less noisy curves. Hence we could estimate the opening velocity on a much smaller timescale, looking at the first 250ms. Note that the gradation observed is still found if we fit curves independently on a longer timescale (as it is done in the notum), and then average velocities for each junction type. This strategy was actually the one used in our previous paper (Chan et al. 2017), and yielded a similar gradation. Velocities determined here are closer to the actual “initial” velocity, as they are measured on a shorter timescale after the cut. The higher values found here suggest that it is indeed the case.

In the germband, the opening velocities were determined using PIV, as several junctions are involved in the opening process. The measurement routine was described in a previous article (Collinet et al. 2015). In short, PIV is computed between a snapshot taken upon ablation and a snapshot taken 2s after ablation. The velocity field is averaged in a region adjacent to the cut line to obtain a scalar velocity value.

---

**Figure S1.**
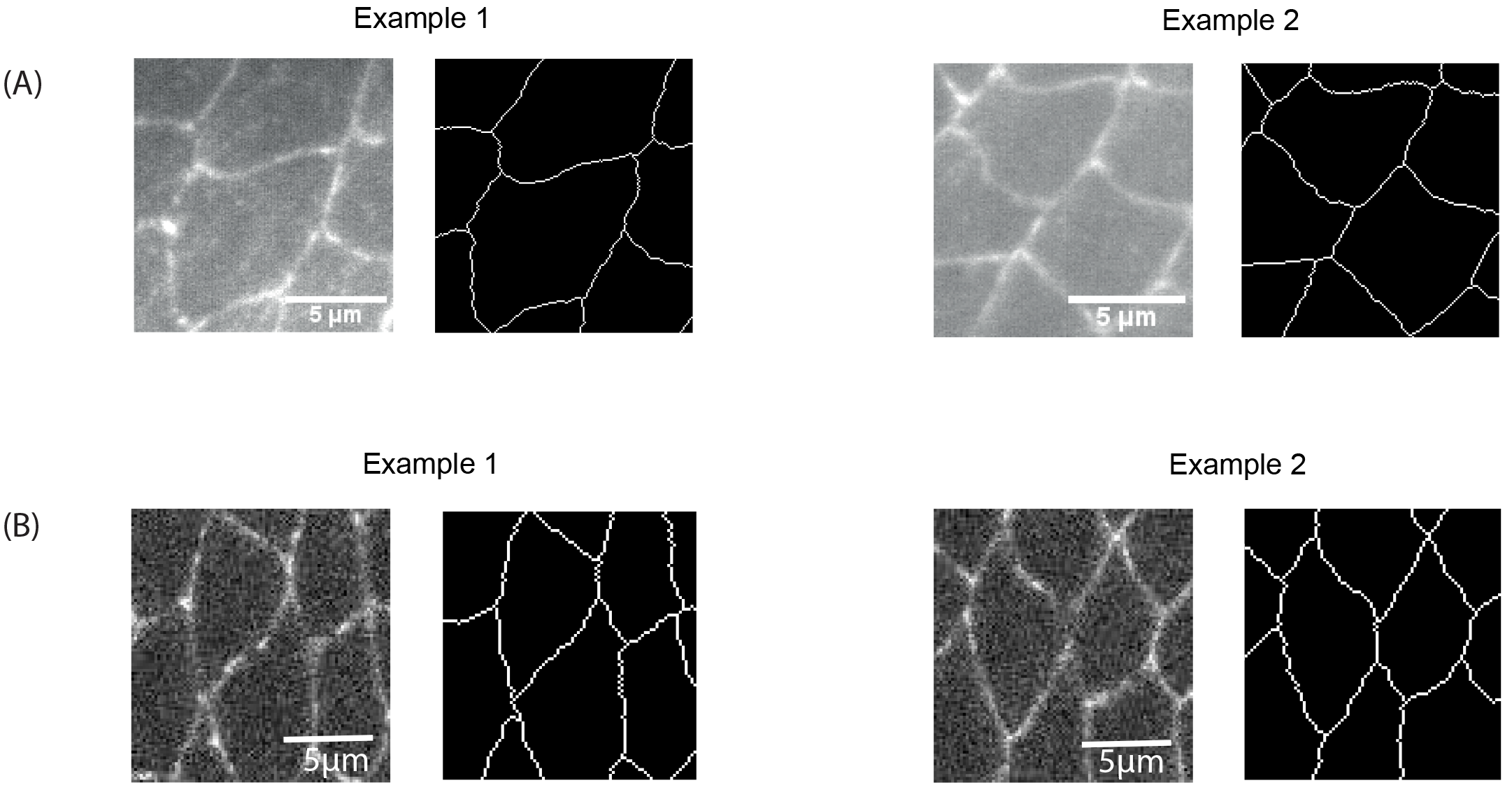
Examples of S shaped junctions. **(A)** Two examples in the notum (left, original image, right, segmented image). **(B)** Two examples in the germband (left, original image, right, segmented image).

**Figure S2.**
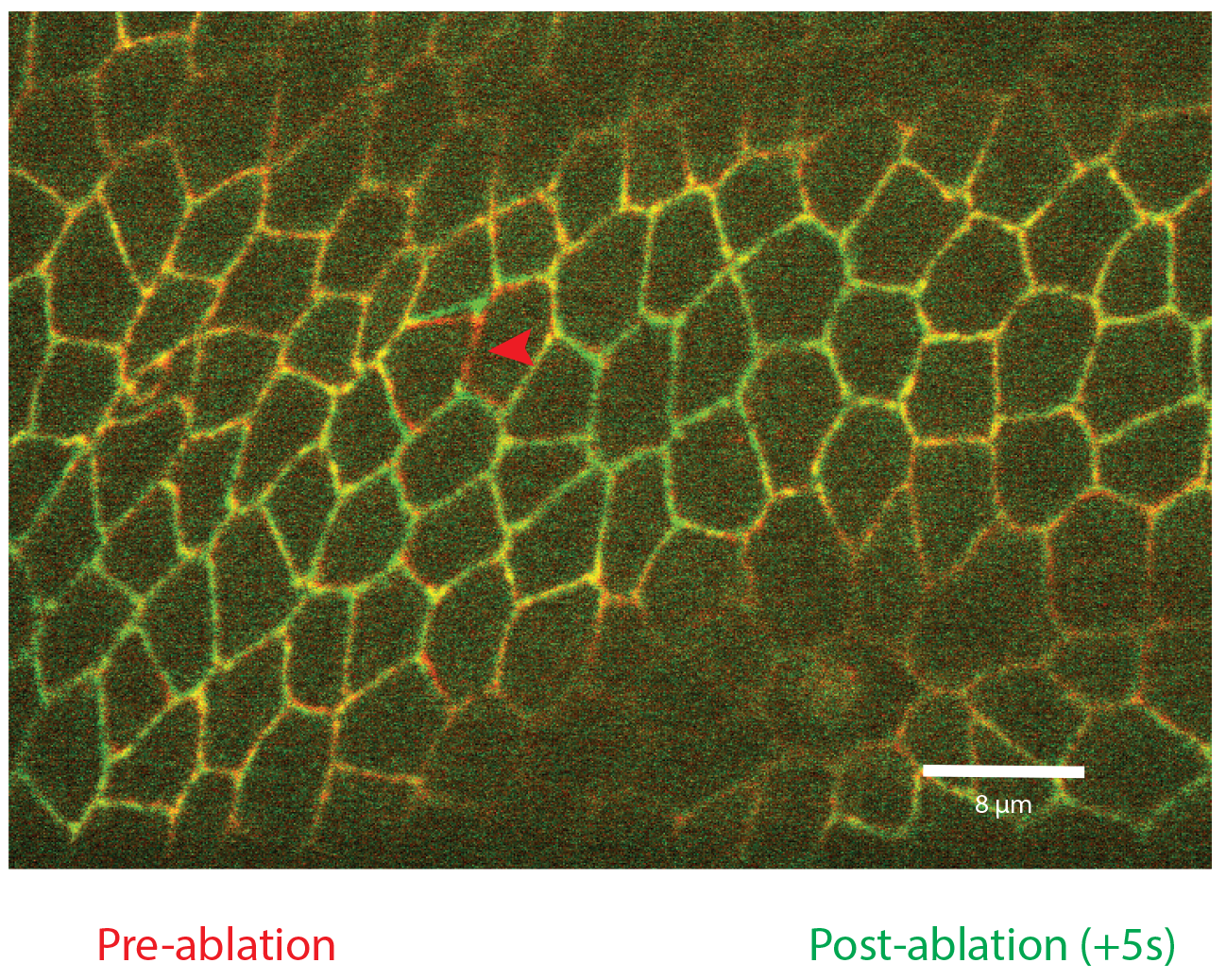
Impact of point ablation on tissue geometry. Overlay of the tissue geometry prior (red) and after (green) cutting a single junction marked by an arrowhead). The displacement in the surrounding cells decays rapidly, and is hardly noticeable above a one-cell distance.

**Figure S3.**
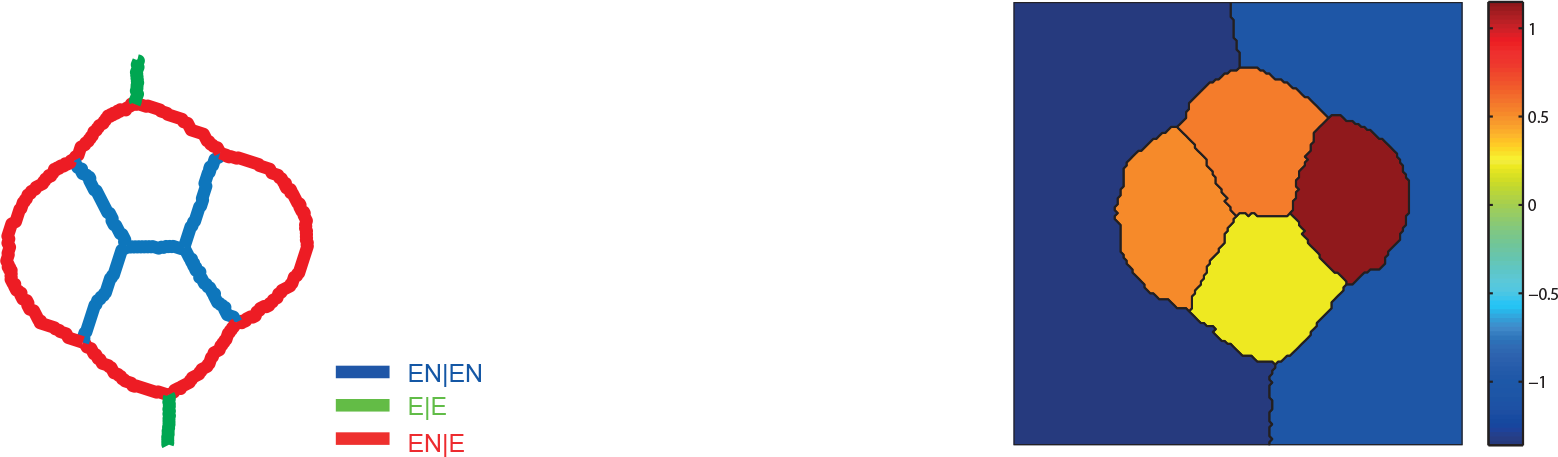
Inferred pressures in the wild type ommatidium. Left: Junction types. Right: Pressure map obtained using Laplace force inference in the averaged wild type ommatidium.

**Figure S4.**
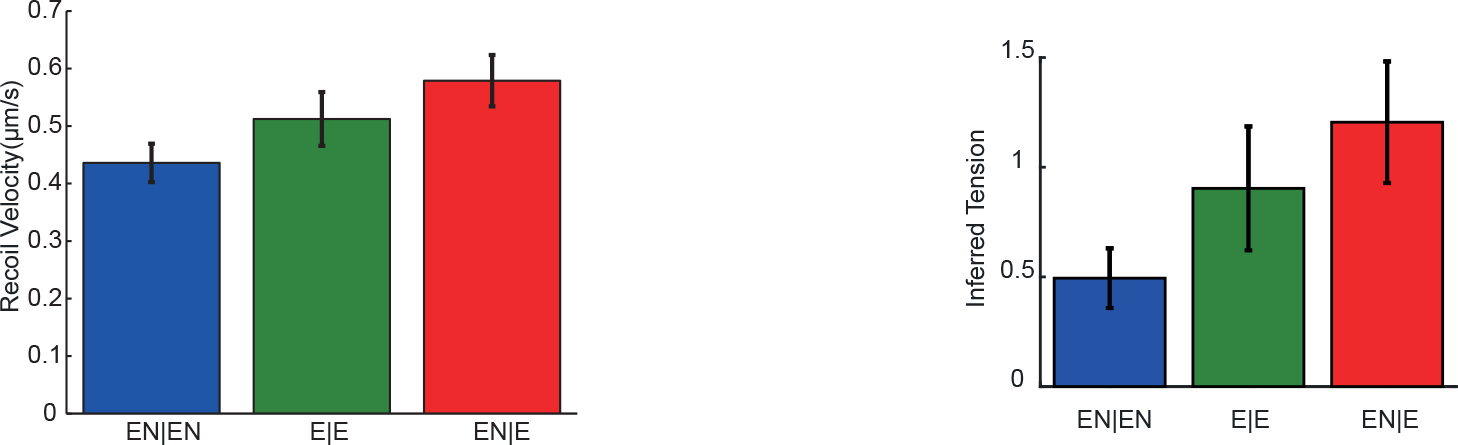
Inferred tensions vs opening velocities in mutant ommatidia. Average opening velocity measured for each junction type, averaged over all mutant configurations (left) and average tension inferred for each junction type, averaged over all mutant configurations (right).

**Figure S5.**
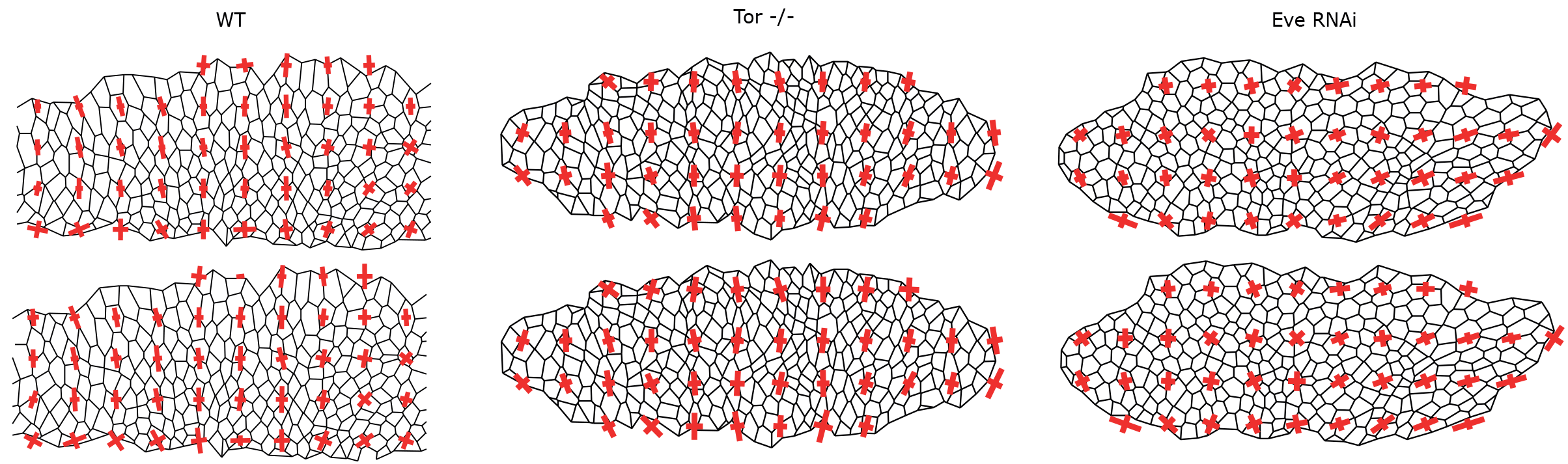
Stress map with and without force inference. Stress maps obtained for each condition using Batchelor’s formula and using tensions and pressures inferred with Bayesian inference (top), compared to the stress maps obtained for each condition using Batchelor’s formula assuming that all tensions are equal to 1 and all pressures are equal to 0 (bottom).

